# The influence of Sex on microRNA expression in Human Skeletal Muscle

**DOI:** 10.1101/2023.02.27.530361

**Authors:** Danielle Hiam, Shanie Landen, Macsue Jacques, Sarah Voisin, Séverine Lamon, Nir Eynon

## Abstract

**Introduction:** Sex differences in microRNA (miRNA) expression profiles have been found across multiple tissues. Skeletal muscle is one of the top tissues that underpin sex-based differences, yet there is limited research into whether there are sex differences in miRNA expression in skeletal muscle. Further, there is limited literature investigating potential differences between males and females in skeletal muscle miRNA expression following exercise, a well-known modulator of miRNA expression. Therefore, the aim of this study was to investigate the effect of sex on miRNA expression in skeletal muscle at baseline and after an acute bout of exercise.

**Methods:** MiRNAs were measured using Taqman®miRNA arrays in skeletal muscle of 42 healthy participants from the GeneSMART study (24 males and 20 females aged 18-45 yrs). Differentially expressed miRNAs were identified using mixed linear models adjusted for age. Experimentally validated miRNA gene targets enriched in skeletal muscle were identified in-silico. Over representation analysis was conducted to identify enriched pathways. TransmiR V.2 was used to identify transcription factor (TF)-miR regulatory networks using CHIP-derived data. We further profiled the effects of two sex-biased miRNAs overexpressed in human primary muscle cells lines derived from male and female donors to understand the transcriptome targeted by these miRNAs and investigate and potential sex-specific effects.

**Results:** A total of 80 miRNAs were differentially expressed in skeletal muscle between the sexes, with 61 miRNAs responding differently to the exercise between the sexes. Sex-biased miRNA gene targets were enriched for muscle-related processes including proliferation and differentiation of the muscle cells and numerous metabolic pathways, suggesting that miRNAs are playing a role in programming sex differences in skeletal muscle. Over-expression of sex-biased miRNAs miRNA-30a and miRNA-30c resulted in profound changes to gene expression profiles that were partly specific to the sex of the cell donor in human primary skeletal muscle cells.

**Conclusion:** We found sex-differences in the expression profile of skeletal muscle miRNAs at baseline and in response to exercise. These miRNAs target regulatory pathways essential to skeletal muscle development and metabolism, suggesting that miRNAs play a profound but highly complex role in programming sex-differences in the skeletal muscle phenotype.

## Introduction

Biological sex is a fundamental characteristic that has profound effects on physiological and molecular factors influencing nearly all human traits [1, 2]. Sex differences arise from a combination of interacting factors such as differences in sex hormone exposure, sex chromosome complement, epigenetic mechanisms and non-coding RNA programming. Whilst sex hormones and the sex chromosomes are the major drivers of sex difference, the importance of non-coding RNAs, specifically microRNAs (miRNAs), in programming sex differences is becoming apparent [3, 4].

MiRNAs are short (~20nt) single-stranded molecules that primarily act by repressing a target mRNA molecule and reducing the translation and expression of the corresponding protein [5]. As regulators of gene expression, they play a crucial role in a variety of biological processes, such as cell development, cell proliferation, differentiation, apoptosis, and cellular signalling. Sex differences in miRNA expression profiles have been found across multiple tissue including pancreatic islets [6], peripheral blood, brain and mucosa tissues [3]. There is however limited research into whether there are sex differences in miRNA expression in skeletal muscle. Yet, skeletal muscle is one of the top tissues that underpin sex-based differences and has the second highest number of genes (up to 3000) that are differentially expressed between the sexes [7, 8]. This indicates a potential role of upstream regulatory mechanisms, such as miRNAs, which could be crucial in programming sex differences in gene expression profiles and the resulting phenotype.

Males and females display distinct muscle phenotypes. On a functional level, females have lower muscle mass and strength compared to males [9, 10]. On the molecular level, there are intrinsic sex differences in fibre type proportions, satellite cell numbers and substrate metabolism [11]. These functional and molecular differences result in females being more susceptible to negative metabolic and functional consequences of age-related muscle loss, as well as prolonged periods of disuse due to sedentary lifestyle or muscle pathologies [9, 11]. Exercise is one of the most potent stimuli to skeletal muscle and is therefore one of the most effective lifestyle management therapies for preventing age-related muscle loss and metabolic conditions such as type 2 diabetes mellitus (T2DM) and cardiovascular disease [12]. Males and females display differences in musculoskeletal, cardiovascular and metabolic responses to the same exercise regime [13]. These differences have been partially attributed to differences in sex hormones [14, 15], transcription factors [16] and more recently DNA methylation [17, 18]. Exercise is well known to modulate microRNA expression [19], but there is limited literature investigating potential differences between males and females in skeletal muscle miRNA expression following exercise. Therefore, the aim of this study was to investigate the role of sex in miRNA expression in skeletal muscle at baseline and after an acute bout of exercise.

## Methods

### Participants

The tissue used in this study was from the Gene and Skeletal Muscle Adaptive Response to Training (Gene SMART) cohort, which is a part of on-going biobank[20]. The detailed methodology has been previously published [21–23]. Briefly, 24 healthy males (age = 32 ± 8.1 years-old; BMI = 25.0 ± 3.4 kg/m2), and 20 healthy pre-menopausal females (age = 34.6 ± 7.4 years-old; BMI = 23.4 ± 2.9 kg/m2) participated in the study. This study was approved by the Human Ethics Research Committee at Victoria University and all participants provided written informed consent.

### Aerobic Capacity (Graded exercise test)

Aerobic capacity was assessed by a graded exercise test (GXT) performed on an electronically braked cycle-ergometer (Lode-Excalibur sport, Groningen, the Netherlands) to measure maximal oxygen uptake 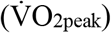 and peak power output (W_peak_). The 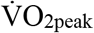 was determined using a calibrated Quark CPET metabolic system (COSMED, Rome, Italy). The GXT consisted of four-minute stages separated by 30 second rest periods until voluntary exhaustion with incremental increases in resistance at each stage. Capillary blood samples were collected at the end of each four-minute stage and immediately after exhaustion and were analysed by the YSI 2300 STAT Plus system (Ohio, USA) to establish lactate concentration. Lactate Threshold (LT) was calculated by the modified DMAX method as previously described [21]. The GXT was performed in duplicate at both baseline and after the intervention and the average was calculated for all parameters between the two tests. In addition, at baseline, participants performed a familiarisation test of the GXT.

### Diet control (48h prior to testing)

To standardise diet across the participants and minimise the effects of this confounding factor, each participant was provided with an individualised pre-packaged diet 48 hours prior to providing the blood samples [24, 25]. The energy content of the provided meals was calculated using the Mifflin St-Jeor equation using the participant’s body mass, height and age [26]. The content of the diets was based on the current Australian National Health and Medical Research Council (NHMRC) guidelines. Participants were asked to abstain from caffeine and alcohol throughout the 48-hour diet as well as food consumption 12 hours prior to blood and muscle collection.

### Blood collection

Blood samples were collected at rest and three hours after the acute bout of HIIE. Venous blood samples were collected via venepuncture or cannulation in BD SST Vacutainers (Becton and Dickson Company, USA). They were left at room temperature (10 mins) before being centrifuged at 3500 rpm for 10mins at 4°C. Serum was collected and stored at −80°C.

### Muscle collection

Muscle biopsies were taken from the participants vastus lateralis muscle under local anaesthesia (1% xylocaine) using a Bergström needle modified to include suction [27] at baseline and three hours after the acute exercise bout. The muscle was immediately frozen in liquid nitrogen and stored at −80 °C for subsequent analyses. A small piece of the muscle (~10-15mg) was placed in serum free Ham’s F-10 medium (Life Technologies) for primary muscle cell culture.

### Hormone analysis

As the age of our cohort was 18-45 yrs, we measured the two major circulating sex hormones for this life stage, testosterone, and E2 [28, 29]. The assays were completed in the accredited pathology laboratory at Monash Health, Australia. Testosterone was measured using high performance liquid chromatography–mass spectrometry (HPLCMS/MS) method using a liquid sample extraction (AB Sciex Triple Quad 5500 LC/MS/MS system). Estradiol (E2) was measured using a competitive binding immunoenzymatic assay performed on a Beckman Coulter Unicel DXI 800 analyser.

### Acute HIIE bout

Male and female participants completed HIIE on an electronically braked cycle ergometer (Velotron, Racer Mate Inc, Seattle, USA). Participants completed approximately five minutes of warm up at an intensity of their own choosing [range 25-60W] and then cycled for six x two-minute intervals and this was interspersed with 1-min recovery periods at a power of 60 W (work to-rest ratio of 2:1). Intensity was individually determined based on baseline GXT results and calculated as power at lactate threshold (LT) + 40% of the difference between peak aerobic power (W_peak_) and power at LT.

### MicroRNA Profiling of skeletal muscle

RNA was extracted with Tri-Reagent Solution (Ambion) as per manufacturer’s protocol. RNA concentrations (243.3ng/μl ± 81.7ng/μl) and quality (A260/280 ratio = 1.97 ± 0.07) were determined by using the Nanodrop 1000 Spectrophotometer (Thermo Fisher Scientific). A total of 360ng of RNA was reverse transcribed as using Taqman microRNA RT kit and Megaplex RT primers, Human Pool A and B v3.0 (Life Technologies, Australia) as per manufacturer’s protocol. MiRNA expression was measured using the TaqMan array, Human microRNA A and B (Life Technologies, Australia). Collectively, these cards allow for the accurate quantitation of 758 human miRNAs. The data was first normalised using global normalisation in cloud based ThermoFisher software which applies a constant scaling factor to every miRNA, so they all have similar median intensity minimising batch effect (supplementary figure 1). The following conditions were used to remove miRNAs that were not expressed in the skeletal muscle for analysis: miRNAs with an average Cq greater than 35 across all samples were excluded (n = 105), miRNAs where more than 20% of the samples (18/88) had a Cq of 35 were excluded (n = 345). The remaining 308 miRNAs were used in the analysis. The Cq values were then transformed into arbitrary units (AU) using the following equation: AU = (1/2)^Ct^10^10^. The cut-off for the relevant level of expression of each miRNA was set at a mean (Ct) < 35, as recommended by the manufacturer. Out of the 758 miRNAs measured, 450 (58%) were not expressed in skeletal muscle and were excluded from further analysis. The remaining 308 miRNAs were used in the analysis.

### Human Primary Cell Lines

#### Experiment Overview

Human primary cell lines were established from muscle biopsies of three females (Age: 35.3 ± 8.6yrs) and three males (32.3 ± 11.6yrs) from the GeneSMART cohort. A small piece of the muscle biopsy (please see above for detailed biopsy method) was collected. Satellite cells were isolated by dissociation with 0.05% trypsin/EDTA (Life Technologies). Cells were plated on a flask coated with extracellular matrix (ECM; Sigma-Aldrich, Castle Hill, Australia) and allowed to proliferate to 70% confluence before passaging. Myoblasts were maintained in proliferation media [Ham’s F-10 medium (Life Technologies) containing 20% FBS, 25 ng/mL fibroblast growth factor (bFGF; Promega), 1% penicillin streptomycin, and 0.5% amphoteromycin (Life Technologies)] at 37°C and 5% CO2. Cells were passaged by mechanical disturbance twice and then frozen down in proliferation media and 10% DMSO. Upon thawing, cells were passaged once before myoblasts were purified using CD56+ Microbeads (Miltenyi Biotec) to eliminate fibroblasts and other cell populations as per manufacturer instructions. At 70-80% confluence cells were seeded into 6 well plates and were allowed to differentiate by replacing proliferation media to differentiation media [DMEM (Life Technologies) containing 2% HS (New Zealand origin; Gibco, Life Technologies) and 1% penicillin/streptomycin. Medium was replaced every 48hours. On day five of differentiation, myotubes were transfected with either 20nM of miR-30a or miR-30c mimics or 20 nM of a mimic scramble (*mirVana*™ miRNA mimics, ThermoFisher) using Lipofectamine® RNAiMAX Transfection Reagent (Ambion, Life Technologies) diluted 16.7X in Opti-MEM I Reduced serum medium (Life Technologies). This solution was added to differentiation medium at a 10x dilution. After 24 h transfection cells were harvested for RNA [30].

#### RNA extraction and RNA sequencing

RNA was extracted from human primary cells using AllPrep RNA/RNA/miRNA universal kit (Qiagen) according to manufacturer instructions. RNA quantity and quality was established using the Agilent Tape Station (Agilent) the average sample yield was 167ng/μL ± 76.8ng/μL and the RIN average was 9.6 ± 0.2. The RNAseq libraries were prepared using the Illumina TruSeq Stranded Total RNA with Ribo-Zero Gold protocol and sequenced with 150-bp paired-end reads on the Illumina Novaseq6000 (Macrogen Oceania Platform). Reads underwent quality check with FastQC (v0.11.9); Kallisto (v0.46.1) was used to map reads to the human reference genome (*HomoSapien GRCh38*) and to generate transcript counts. Differential gene analysis was conducted using the R package DeSeq2 (REF) on genes with >1 reads per million (RPM) per sample on average. Differentially expressed genes were considered significant at FDR < 0.1, unless otherwise specified.

#### Statistical and Bioinformatic Analysis

All data were analysed using R studio 4.1.3 [31]. Missing data were imputed using the mice package [32] and iteration three was randomly selected for all analyses. To identify differences in miRNA expression between sexes and after exercise two different linear mixed models were used as implemented in the *variancePartition* package in R [33]. Participant ID was used as the random effect to account for repeated measures and allow each individual to have their own intercept. All models were adjusted for age. The first model was conducted in each sex separately to identify what miRNAs were changed three hours after an acute bout of exercise. The model was of the form: miRNA ~ Timepoint + Age + (1|ID). The second model was used to tease out differences between sex regardless of the timepoint, and the interaction was used to determine the differential response of miRNAs to an acute bout of exercise between sexes. The model was of the form: miRNA ~ Sex * Timepoint + Age + (1|ID).

Hormone analysis was of the form miRNA ~ Hormone + (1|ID) and, due to collinearity, were run separately for each hormone and sex. A chi square test was performed to investigate if there was an over- or under-representation of sex-biased miRNAs on a specific chromosome. Gene target analysis was conducted using the R package *multi-MiR* [34]. Only gene targets that had been experimentally validated were included in the gene target and pathway enrichment analysis. Over representation analysis (ORA) was used for gene target enrichment and implemented by the R package *clusterProfiler* [35]. The background gene list used for the ORA analysis comprised all experimentally validated targets of the miRNAs detected in the microarray. Tissue enrichment of the validated gene targets was conducted in the R package *TissueEnrich[36].* Finally, transcription factor-microRNA interactions were explored using *TransmiR v2.0* [37].

Benjamini-Hochberg adjustment was used to correct for multiple comparisons to minimize the risk of false positive results and P values with an FDR <0.05 were deemed statistically significant unless otherwise stated. Data are presented as Mean ± SD unless stated otherwise. The following packages were also used in our analysis; *lme4* [38], *lmerTest* [39], and *tidyverse* [40]. The full R code can be found at https://github.com/DaniHiam/SexBiasedmiRNAs.

## Results

There were no differences in age, body mass index (BMI), aerobic capacity (VO_2peak_) or lactate threshold (LT) between males and females. Males had a higher W_peak_ than females. As expected, there were significant differences in circulating sex hormones between males and females (Table 1).

**Table 1:** Participant characteristics stratified by Sex.

|  | Female | Male | p |
| --- | --- | --- | --- |
| *n* | 20 | 24 |  |
| *Age (years)* | 34.61 (7.37) | 31.96 (8.14) | 0.268 |
| *BMI (kg.m<sup>-2</sup>)* | 23.45 (2.90) | 24.98 (3.42) | 0.122 |
| *W<sub>peak</sub> (W.kg<sup>-1</sup>)* | 3.18 (0.71) | 3.90 (0.88) | 0.005 |
| *VO<sub>2</sub> (mL.kg<sup>-1</sup>min<sup>-1</sup>)* | 42.81 (7.08) | 47.21 (11.91) | 0.155 |
| *LT (W.kg<sup>-1</sup>)* | 2.14 (0.61) | 3.85 (5.41) | 0.169 |
| **Circulating sex hormones** |  |  |  |
| *TT (pmol/L)* | 0.84 (0.29) | 19.61 (5.11) | <0.001 |
| *fT (pmol/L)* | 9.96 (3.92) | 473.63 (140.71) | <0.001 |
| *SHBG (nmol.L)* | 73.11 (33.60) | 34.87 (15.44) | <0.001 |
| *Estrogen (pmol/L)* | 227.55 (217.01) | 107.73 (24.66) | 0.012 |

BMI, Body Mass Index; W, watts; LT, Lactate Threshold; TT, Total Testosterone; fT, Free Testosterone; SHBG, Sex Hormone Binding Globulin. Results are presented as mean (SD). P values <0.05 are considered significant.

### miRNA Results

#### MiRNA expression at baseline and following exercise

In the female cohort, seven miRNAs were upregulated in response to an acute bout of exercise, while in the male cohort, eight miRNAs were downregulated, and 113 miRNAs were upregulated in response to an acute bout of exercise (Fig 1A and supplementary file 1). Only one miRNA, miR-30d-5p, overlapped between sexes, however in females it was up-regulated with exercise while in males it was down-regulated with exercise (Fig 1B). A total of 80 miRNAs were differentially expressed between the sexes regardless of the timepoint. MiR-22-5p displayed a higher expression in males compared to females, while the remaining 79 miRNAs displaying a lower expression in males compared to females (Fig 1C). Interaction analysis indicated that 61 miRNAs responded differently to the exercise between the sexes. Nine miRNAs were more highly expressed in females after an acute bout of exercise compared to males. The remaining 52 miRNAs were more highly expressed in males after an acute bout of exercise compared to females (Fig 1D).

**Figure 1:**
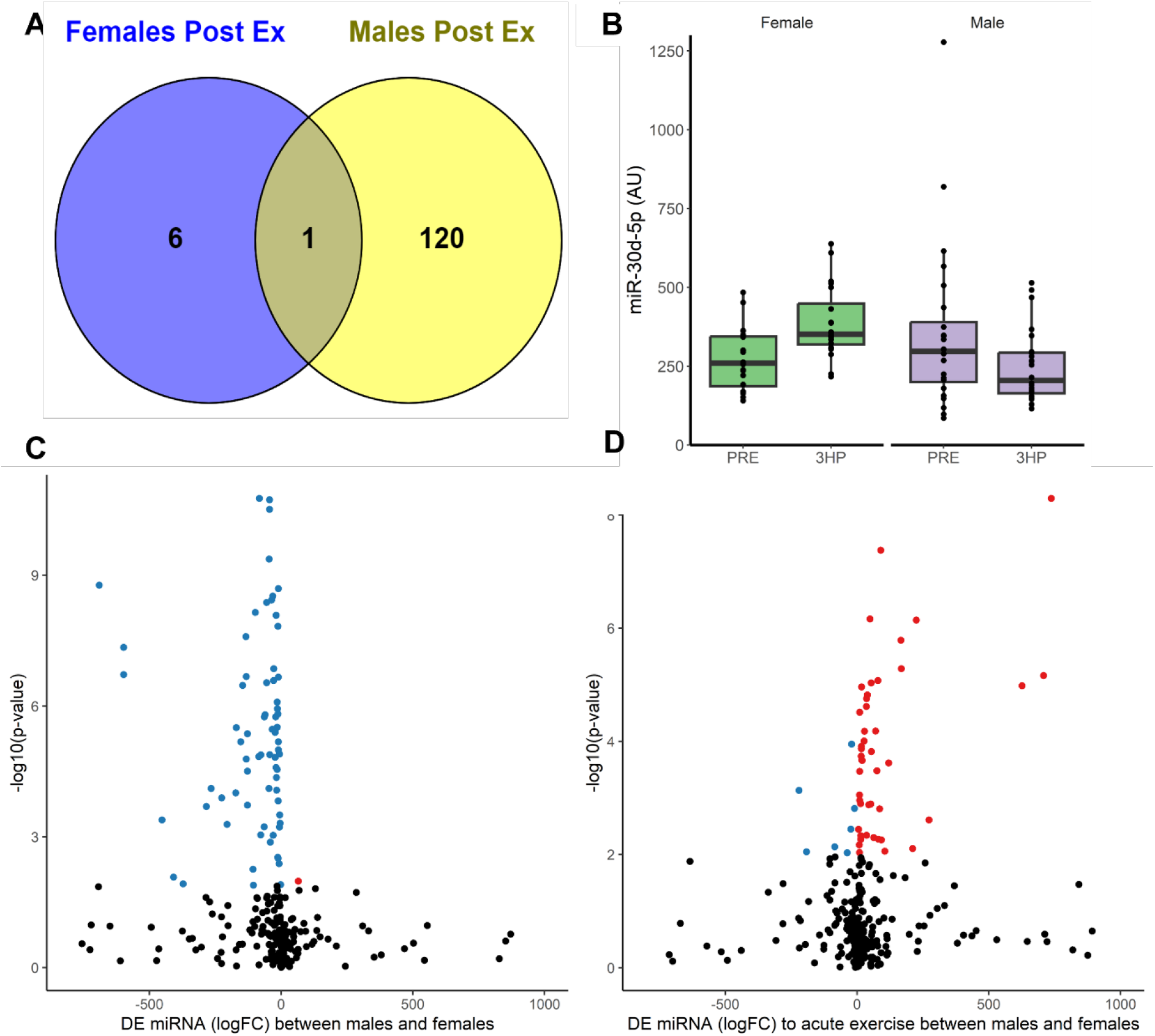
The skeletal muscle microRNA profile displays profound differences at baseline and in response to exercise between males and females. **A)** Venn Diagram displaying number of miRNAs that were altered 3 hours after an acute bout of exercise in females (Females Post Ex) and males (Males Post Ex); FDR <0.1. **B)** Relative expression of miR-30d-5p at pre (PRE) and three hours post-acute exercise (3HP) faceted by sex **C)** Volcano Plot: differences in microRNA expression levels between males compared to female. Each point represents microRNA, red points indicated an increase in miRNA expression in males compared to females. Blue points indicated a decrease in miRNA expression in males compared to females. Black dots were not significant. **D)** Volcano Plot: Sex-specific differences in miRNA expression in response to an acute bout of exercise.

#### Over-representation pathway analysis

Over-representation analysis (ORA) was performed to understand the gene pathways targeted by the sex-biased miRNAs. First, we separately investigated the 7 and 121 miRNAs that were altered by exercise in female and males, respectively. In females, 320 Gene Ontology (GO) terms belonging to the ‘biological processes’ category were overrepresented, while in males 15 GO terms of the ‘biological processes’ category were overrepresented (Supplementary File 1). In females, these terms broadly related to “maintenance of cell number”, “extracellular vesicle biogenesis”, “circadian regulation of gene expression” and “centrosome localisation”. In males, they broadly related to “proteasome-mediated ubiquitin-dependent protein process, “tissue migration”, “mitotic cell cycle transition” and “ribonucleic complex biogenesis” (Figure 2A).

**Figure 2:**
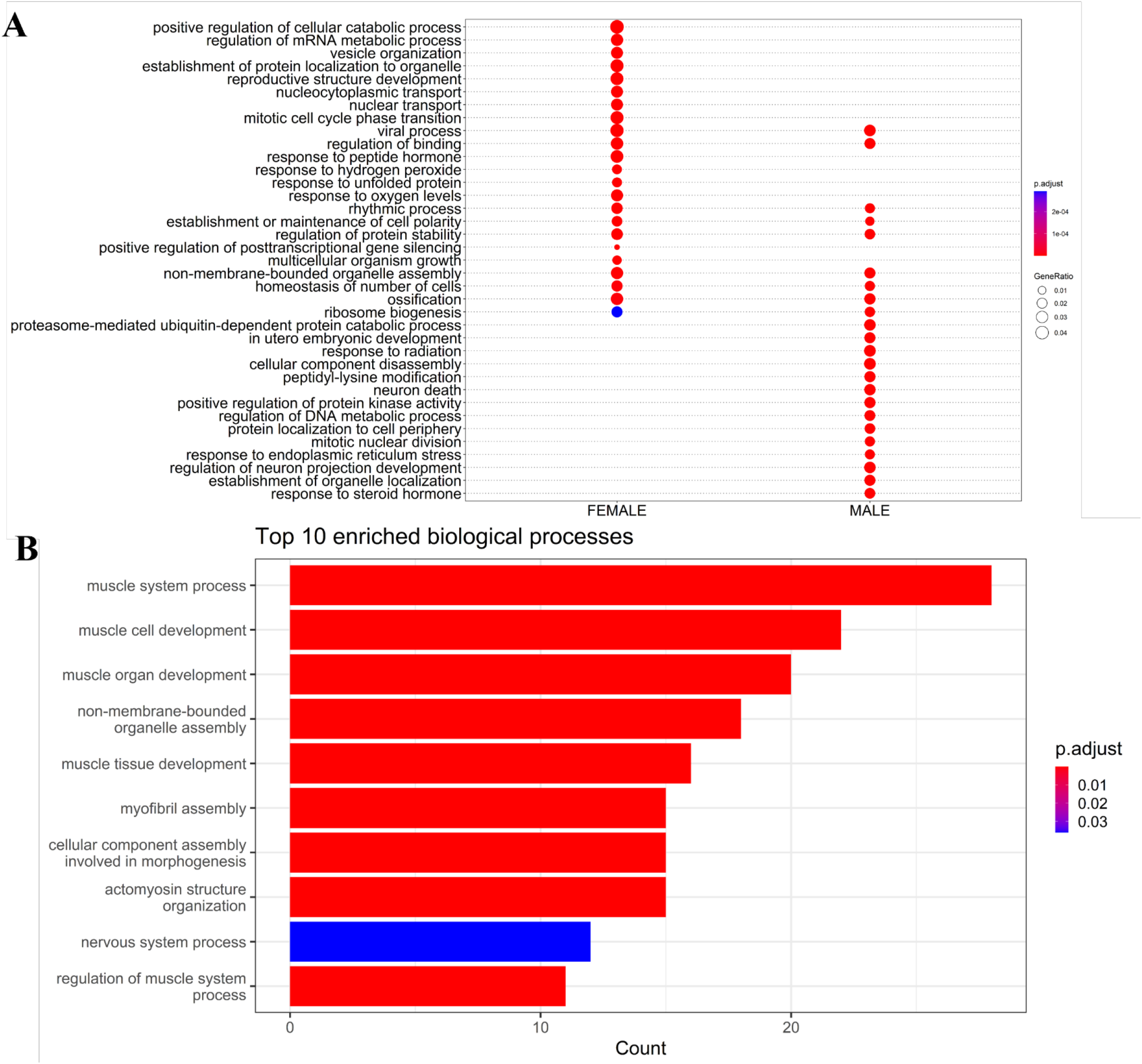
Gene set enrichment analysis of the gene targets of sex-biased miRNAs. **A)** Comparison of the significant gene ontology (GO) terms belonging to the ‘biological processes’ category among males compared to females. **B)** Top enriched GO terms for skeletal muscle enriched gene targets of the sex-biased miRNAs.

We then investigated the overrepresented gene pathways that were putatively targeted by the miRNAs displaying a different response to exercise between the sexes. This analysis was conducted on genes that were considered expressed in skeletal muscle based on RNA sequencing data from the GTEx portal [36]. Overall, 83 Gene Ontology (GO) terms were over-represented in this analysis (Figure 2D, Supplementary File 1). The GO pathways broadly belonged to the ‘biological processes’ categories including “muscle structure development”, “muscle contraction”, “regulation of system processes” and “actomyosin structure organisation” (Figure 2B).

#### Chromosome location of miRNA genes

A fisher test was performed to investigate if sex-biased miRNA genes were more likely to be encoded on the sex chromosomes compared to the autosomes. There was no difference in likelihood of the miRNA gene being located on the sex chromosomes and autosomes (Figure 3A). We then investigated whether miRNAs were more likely to be sex-biased if they were located on a specific chromosome. At baseline, there was an increased chance of a miRNA being differentially expressed by sex if located on chromosome 20 (p=0.02) (Fig 3B). After the acute exercise bout, there was no enrichment of sex-biased miRNAs genes on any specific chromosome (p>0.05).

**Figure 3:**
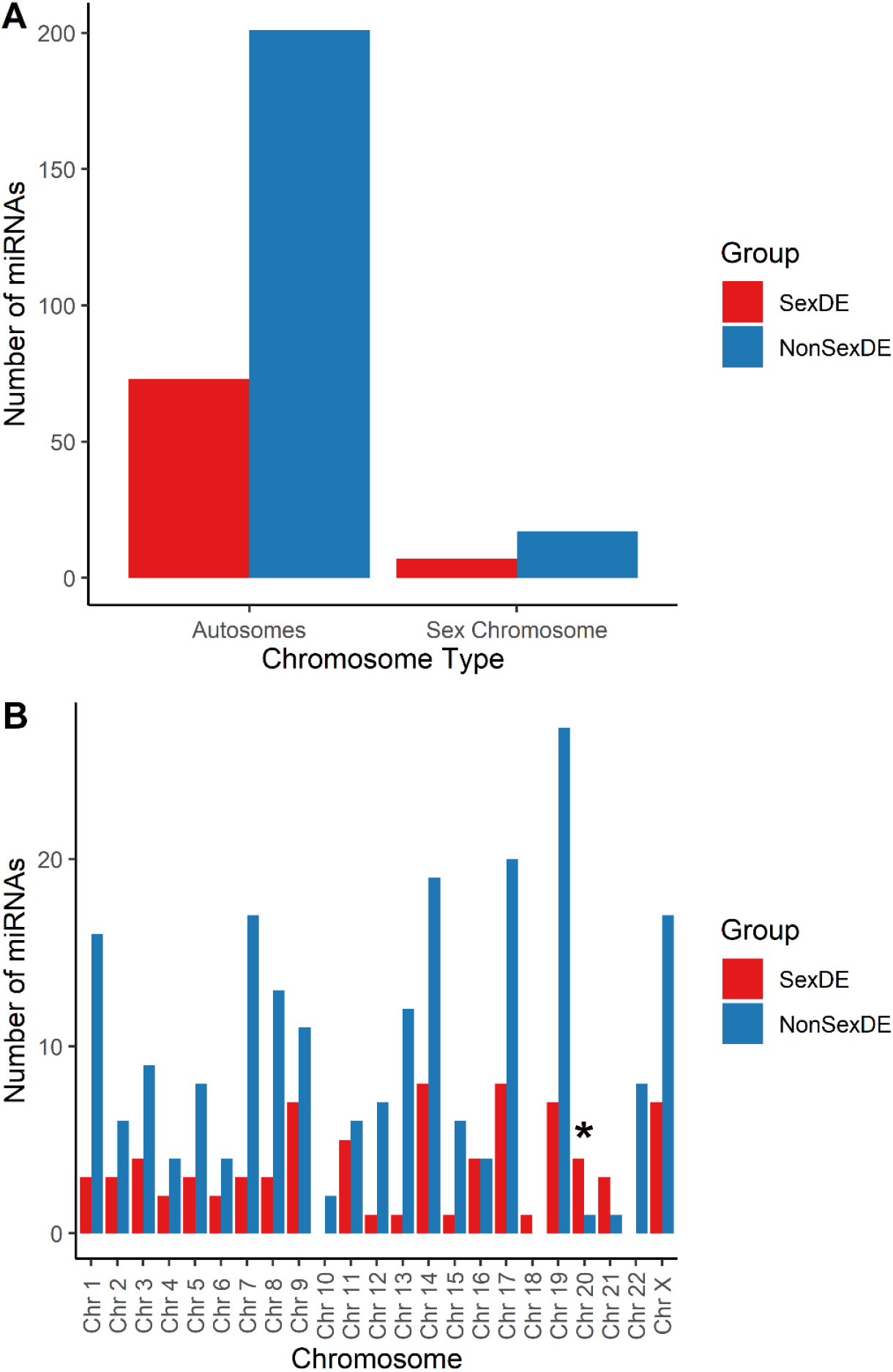
The number and location of the microRNAs expressed across the chromosomes in skeletal muscle. Red indicates the number of microRNAs differentially expressed between males and females (SexDE). Blue indicates the number of expressed miRNAs that did not differ between males and females (NonSexDE).

#### Transcription Factors

We then examined whether there was an enrichment for TFs binding sites located near the transcription start site of sex-biased miRNAs at baseline and in response to exercise. At baseline, the transcription binding sites for AKT1, SMAD3, TGFB1, DNMT1, EPAS1, NFE2L2 and TP53 were enriched. The transcription regions for sex-biased miRNAs in response to exercise were enriched for the binding sites of SMAD3, NFKB1 and TGFB1.

#### Circulating Sex Hormones

To understand the contribution of circulating sex hormone concentration to sex-biased miRNAs, we first investigated miRNAs with a sex hormone response element (HRE) (AR, ESR1 and ERS2) in their transcription start site (TSS). The putative transcriptional regulatory region was considered 300bp upstream and 100bp downstream of each miR TSS [37]. Total testosterone (TT) and free testosterone (fT) were positively associated with miR-20a-3p prior to FDR adjustment (TT: p = 0.03; FDR = 0.66; fT: p = 0.049; FDR = 0.94) in females, but not males (TT: p = 0.09; FDR = 0.99, fT: p = 0.09; FDR = 0.99). Estrogen (E2) was negatively associated with miR-18b-5p (p = 0.002, FDR = 0.07) prior to FDR adjustment in females only.

Sex hormones can also signal indirectly through protein-protein interaction with other TFs [41]. Therefore, we then investigated all miRNAs expressed in skeletal muscle regardless of whether they had a HRE for sex hormones in the TSS (Table 2). TT was associated with 6 miRNAs in females, but only one miRNA in males. FT was significantly associated with 3 miRNAs in females and 11 miRNAs in males. E2 was associated with 17 miRNAs in females, and 8 miRNAs in males. There was no overlap between the miRNAs that were associated with E2, TT or fT between males or females.

**Table 2:** miRNAs associated with circulating sex hormone concentration.

| *Sex Hormone* | FEMALE |  |  | MALE |  |  |
| --- | --- | --- | --- | --- | --- | --- |
|  | miRNA | pVal | adj pVal | miRNA | pVal | adj pVal |
| *TT* | miR-1247-5p | 0.03 | 0.95 | miR-193b-5p | 0.04 | 1.00 |
|  | miR-20a-3p | 0.03 | 0.95 |  |  |  |
|  | miR-550a-5p | 0.03 | 0.95 |  |  |  |
|  | miR-939-5p | 0.04 | 0.95 |  |  |  |
|  | miR-126-5p | 0.04 | 0.95 |  |  |  |
|  | miR-628-3p | 0.05 | 0.95 |  |  |  |
| *fT* | miR-452-5p | 0.03 | 0.99 | miR-132-3p | 0.01 | 0.98 |
|  | miR-939-5p | 0.03 | 0.99 | miR-139-5p | 0.02 | 0.98 |
|  | miR-20a-3p | 0.05 | 0.99 | miR-126-3p | 0.02 | 0.98 |
|  |  |  |  | miR-613 | 0.02 | 0.98 |
|  |  |  |  | miR-522-3p | 0.03 | 0.98 |
|  |  |  |  | miR-339-5p | 0.04 | 0.98 |
|  |  |  |  | miR-491-5p | 0.04 | 0.98 |
|  |  |  |  | miR-10a-5p | 0.04 | 0.98 |
|  |  |  |  | miR-210-3p | 0.04 | 0.98 |
|  |  |  |  | miR-125a-5p | 0.04 | 0.98 |
|  |  |  |  | miR-126-5p | 0.05 | 0.98 |
| *E2* | miR-18b-5p | 0.00 | 0.51 | miR-193b-5p | 0.00 | 0.95 |
|  | miR-15a-5p | 0.01 | 0.82 | miR-1260a | 0.01 | 0.95 |
|  | miR-494-3p | 0.01 | 0.82 | miR-650 | 0.02 | 0.95 |
|  | miR-98-5p | 0.02 | 0.82 | miR-643 | 0.02 | 0.95 |
|  | miR-502-3p | 0.03 | 0.82 | miR-590-3p | 0.03 | 0.95 |
|  | miR-124-3p | 0.03 | 0.82 | miR-519b-3p | 0.04 | 0.95 |
|  | miR-639 | 0.03 | 0.82 | miR-24-2-5p | 0.04 | 0.95 |
|  | miR-148b-3p | 0.03 | 0.82 | miR-1225-3p | 0.04 | 0.95 |
|  | miR-520c-3p | 0.04 | 0.82 |  |  |  |
|  | miR-597-5p | 0.04 | 0.82 |  |  |  |
|  | miR-324-3p | 0.04 | 0.82 |  |  |  |
|  | miR-1253 | 0.04 | 0.82 |  |  |  |
|  | miR-30d-3p | 0.04 | 0.82 |  |  |  |
|  | let-7f-5p | 0.04 | 0.82 |  |  |  |
|  | miR-571 | 0.04 | 0.82 |  |  |  |
|  | miR-1290 | 0.05 | 0.84 |  |  |  |
|  | miR-487b-3p | 0.05 | 0.84 |  |  |  |

pVal, p value; adj, adjusted, TT, total testosterone; fT, free testosterone; E2, estradiol.

#### Overexpression of selected sex-biased miRNAs

All 5 miR-30 family members (miR-30a-5p, miR-30b-5p, miR-30c-5p and miR-30d-5p/-3p) displayed differences in expression between sexes and a sex-specific response to exercise. Some research indicates that the miR-30 family members have a regulatory role in the differentiation of human skeletal muscle cells [42, 43]. MiR-30a-5p and miR-30c-5p have yet to be extensively validated in skeletal muscle. They displayed large differences (logFC > 1.5) in expression between sexes at rest, and a sex-specific response to exercise. Specifically, miR-30a-5p was upregulated after exercise in females but remained unchanged in males and miR-30c-5p which was upregulated after exercise in males and remained unchanged in females. Therefore, we selected these two candidates and overexpressed them in primary skeletal muscle cells to understand their effect on the myocyte transcriptome and to explore putative pathways. Hsa-miRNA-30a-5p (miR-30a) and hsa-miRNA-30c-5p (miR-30c) were overexpressed in three female and three male primary skeletal muscle cultures. Overexpression of miR-30a and miR-30c was confirmed by quantifying transcript levels using qPCR (Fig 4A, B) and led to a 70-fold increase in miR-30a and 80-fold increase in miR-30c.We then conducted RNA sequencing to understand the transcriptome profile changes driven by these miRNAs. At baseline, and without miRNA overexpression, 27 genes were differentially expressed between primary muscle cells cultured from male and female donors(Fig 4C). In total, 247 genes were altered by the over-expression of miR-30a and miR-30c and were represented by the GO pathways broadly belonging to the ‘biological processes’ categories including “protein glycosylation”, “cytosolic transport” and “Golgi vesicle transport” at FDR < 0.2 (Fig 4D).

**Figure 4:**
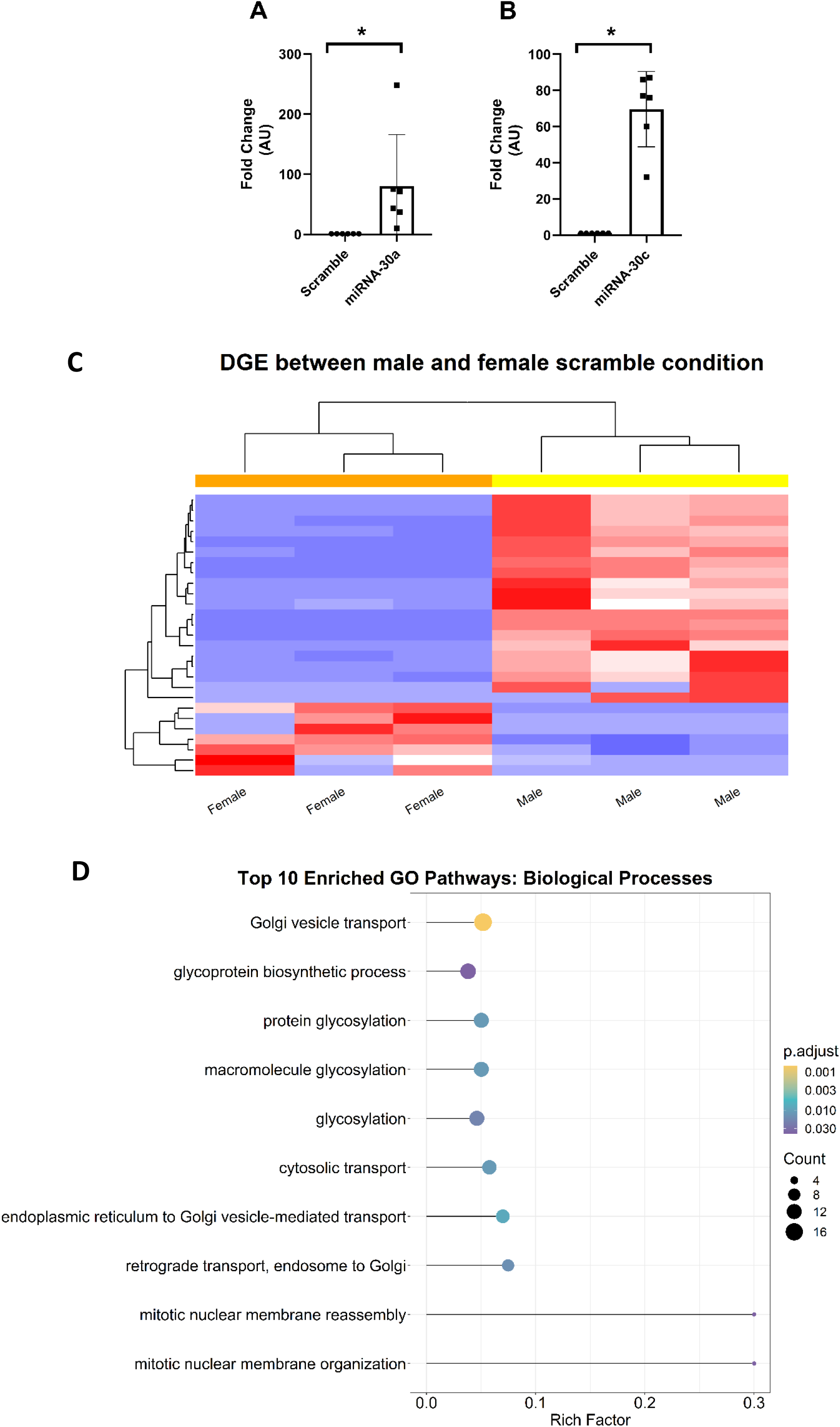
miRNA overexpression in skeletal muscle primary cells. QRT-PCR analysis of miR-30a/30c expression levels containing either **A)** miR-30a or **B)** miR-30c. Fold change normalised back to negative control. *P < 0.05. Data is expressed as Mean ± SD**. C)** Heatmap of top 30 differentially expressed genes clustered by sex **D)** Gene set enrichment analysis: Top enriched GO terms from overexpression of miR-30a and miR-30c. Rich Factor is defined as the ratio of the target genes annotated to a term to the ratio of all genes annotated to that term.

### miRNA-30a

Overexpression of miR-30a resulted in differences in gene expression that were partly specific to the sex of the donor cells. In male-derived primary muscle cell lines, 18 genes were up-regulated, and 30 genes were down-regulated with over-expression of miR-30a (Fig 5A). In female-derived primary muscle cell lines, 18 genes were up-regulated and 33 were down-regulated with over-expression of miR-30a (Fig 5B). Of these, 34 genes were similarly altered between male and female cell lines in response to over-expression of miR-30a. A total of 31 genes were altered differently between male and female cell lines (Fig 5C).

**Figure 5:**
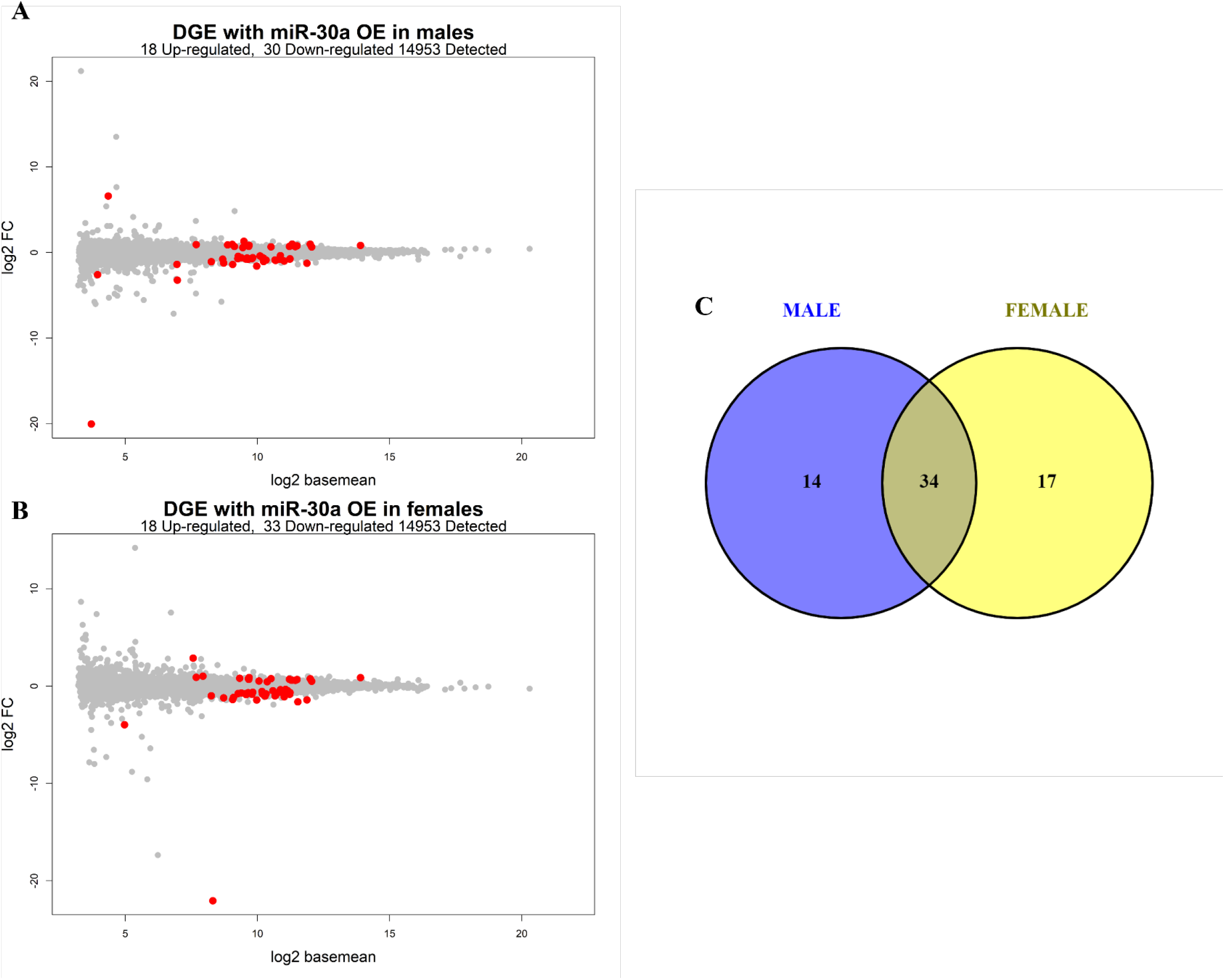
Differentially expressed genes with overexpression of miRNA-30a in human primary skeletal muscle. MA-plot of gene differentially expressed with overexpression of miR-30a in **A)** males or **B)** females. **C)** Venn Diagram comparing the direction of DE genes between the sexes. Significance at FDR < 0.1.

### miRNA-30c

Similar to miR-30a, overexpression of miR-30c resulted in differences in gene expression that were partly specific to the sex of the donor cells. In male-derived primary muscle cell lines, 13 genes were up-regulated, and 14 genes were down-regulated with over-expression of miR-30c (Fig 6A). In female-derived primary muscle cell lines, 12 were up-regulated and 17 were down-regulated with over-expression of miR-30c (Figure 6B). Of these, 11 genes were similarly altered between male and female cell lines in response to over-expression of miR-30c. A total of 34 genes were differently altered between males and females cell lines (Fig 6C).

**Figure 6:**
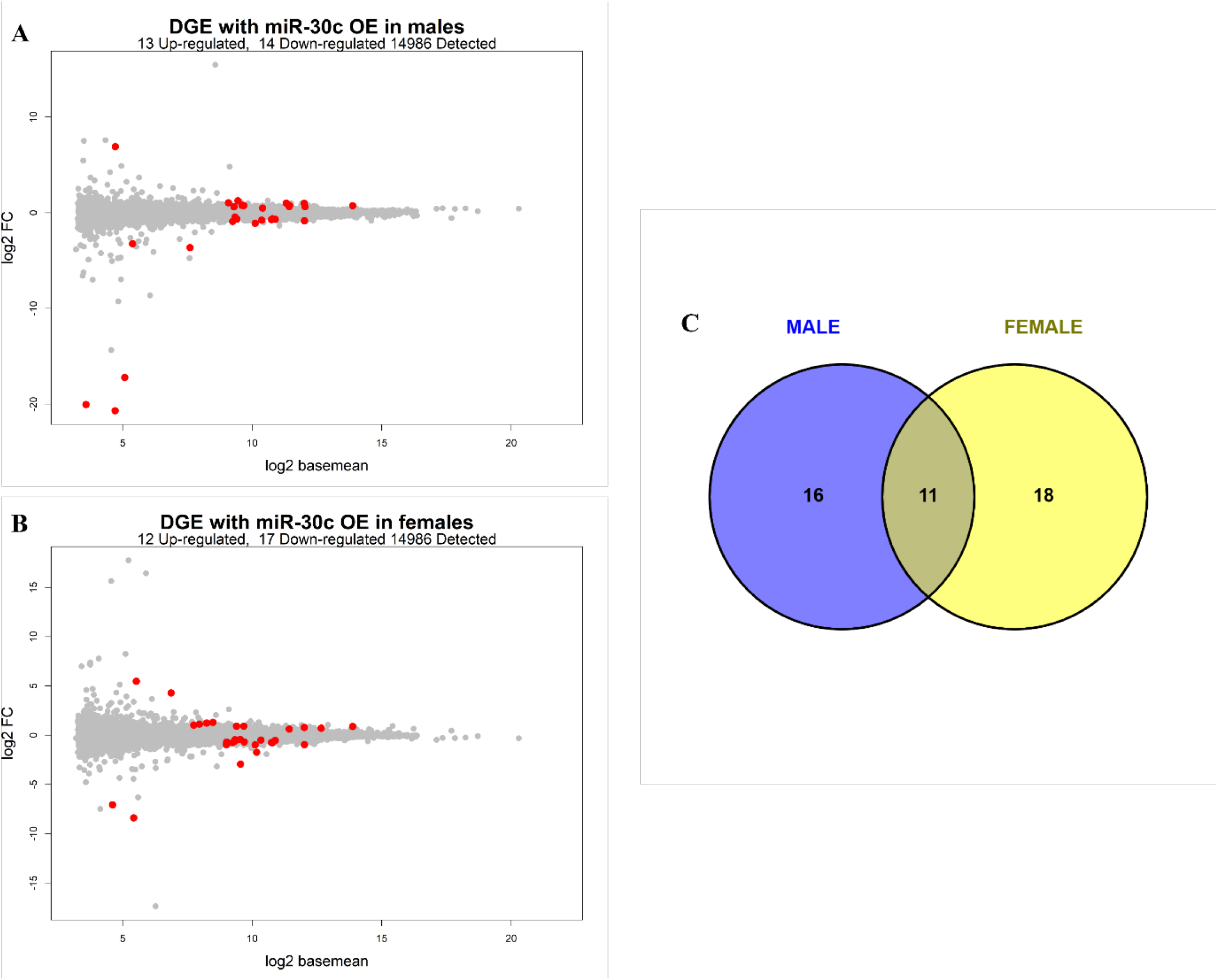
Differentially expressed genes with overexpression of miRNA-30c in human primary skeletal muscle. MA-plot of gene differentially expressed with overexpression of miR-30c in **A)** males or **B)** females. C) Venn Diagram comparing the direction of DE genes between the sexes. Significance at FDR < 0.1.

## Discussion

MiRNAs are necessary for skeletal muscle development [44] and have regulatory roles in determining skeletal muscle phenotype [45], which is one of the most sex-divergent tissues. Few studies have examined the effect of sex on the skeletal muscle miRNAome. Thus, in a screening approach, we conducted miRNA microarrays to profile the skeletal muscle miRNA signature across males and females at baseline and after an acute bout of exercise. We found profound sex differences in the muscle miRNA profile, with an overwhelming number of miRNAs exhibiting a lower expression in males when compared to females. However, three hours after an acute bout of exercise, males exhibited a greater miRNA expression response to an acute bout of exercise compared to females. Sex-biased miRNA gene targets were enriched for muscle-related processes such as proliferation and differentiation of the muscle cells and metabolic pathways, indicating that miRNAs may play a role in programming the physiological sex differences in skeletal muscle pre- and post-exercise. There was no difference in likelihood of the sex-biased miRNA genes being located on the sex chromosomes or autosomes. Multiple miRNAs were associated with sex hormone concentrations, (TT, fT and Estrogen), many of them in a sex-specific way. Finally, the sex-biased miRNAs were enriched for seven transcription factor (TFs) binding sites at baseline and three TFs after an acute bout of exercise indicating the potential role of specific TFs in regulating sex-biased miRNA expression. We further profiled two sex-biased miRNAs to understand the gene expression transcriptome targeted by these miRNAs. Overexpression of miRNA-30a and miRNA-30c in human primary skeletal muscle cells resulted in profound changes to gene expression profiles that were partly specific to the sex of the cell donor.

To our knowledge, this is the most extensive miRNA profiling analysis that has been completed in skeletal muscle comparing males and females. In total, 80 and 61 miRNAs displayed a sex bias at baseline and in response to an acute bout of exercise, respectively. At baseline, an overwhelming majority of miRNAs were more highly expressed in females when compared to males including two myo-miRNAs (muscle-enriched miRNAs), miR-133a/133b. This is in line with previous literature showing that expression levels of miR-133a/133b were markedly higher in females compared to males at baseline [46]. Our findings also suggest that, after an acute bout of exercise, males exhibited a greater change in the miRNA profile compared to females. In males, the myo-miRs-133a/b and 1-3p were upregulated in response to acute exercise in line with most [19, 47, 48] but not all studies [49]. The discrepancies between our data and Silver et.al [49] may be due to the differences in the intensity of the exercise bout between the studies, which can alter miRNA levels differentially [50]. In females, we observed no changes in the expression of miR-133a/b or −1-3p in response to acute exercise, in line with previous findings [49]. MiR-133a/b is under androgenic control in males but not in females, which may explain these sex-specific responses [46]. While miR-133a/b did not associate with testosterone (free or total T) in the male cohort, it is also strongly modulated by aerobic fitness, which may override the regulatory effects of androgens in males [46]. According to Rapp et al.[51], our cohort sat approximately in the 90^th^ percentile for relative VO_2peak_ norms, indicating a high aerobic fitness, which may provide an explanation for this lack of association. Altogether, these data demonstrate the influence of sex on the miRNAome in muscle at both baseline and in response to acute exercise. Future research is required to validate the mechanisms that are programming these sex differences in the miRNA profile of skeletal muscle.

Sex-biased miRNA expression is a result of both hormonal and genetic differences between the sexes. The make-up of sex chromosomes (i.e., XX in biological females and XY in biological males) may contribute to sex differences in miRNA expression as the X chromosome contains a higher density of miRNA genes compared to the Y chromosome. While many miRNAs are inactivated on the X chromosome, others are known to “escape” X inactivation [52]. The current data did not indicate an over-representation of miRNAs on the X chromosome either at baseline or after exercise, but sex-biased miRNAs were more likely to be located on chromosome 20 compared to all other chromosomes. This may be an early indication that sex-biased miRNAs are more likely located on the autosomes than the sex chromosomes, which would be consistent with findings in brain, colorectal tissue, blood and cord blood tissues [3]. Overall, our findings indicate that the sex chromosomes may not play a major role in sex-biased miRNA expression in skeletal muscle, which may rather stem from the differential regulation of miRNA genes located on the autosomes. Further research is required as our findings were limited to miRNAs represented in the microarray array and therefore may not apply to the entire miRNAome of skeletal muscle.

Sex steroid hormones can regulate miRNA expression directly via binding to their nuclear hormone receptors, or indirectly by binding to the host gene promoters in intragenic miRNAs or through protein-protein interactions with other TFs [41]. Our data suggest that the associations between sex hormones and miRNAs expression levels are sex-specific. For example, miRNA-20a-3p showed a positive association with both TT and fT and was upregulated after exercise in females but not males. Limited data is available in skeletal muscle, but other tissues have shown differences in miR-20a-3p expression between sexes. In astrocytes, miRNA-20a-3p is more highly expressed in females compared to males and associated with improved stroke outcomes, which has a sex-biased incidence [53]. The functional significance of miR-20a-3p in skeletal muscle is relatively unknown, however, in one study, it was significantly down-regulated during skeletal muscle cell differentiation in human cells [42]. Further, females with polycystic ovary syndrome (PCOS), characterised by elevated testosterone, displayed a higher miR-20a-3p expression in follicular fluid compared to females without PCOS [54]. In-silico analysis indicates that there is an androgen response element (ARE) located near the miRNA genes’ transcription start site and this may provide an explanation for the positive association between miRNA-20a-3p and androgen concentration. We can only speculate as to why this observation is not also exhibited in males. Previous literature has shown that transcription factors have a sex-biased targeting patterns and indeed skeletal muscle has one of the most sex-divergent TF targeting patterns [16]. Several mechanisms could be mediating these patterns, including TF abundance and conformation as well as epigenetic modifications such as DNA methylation. Our group recently found enrichment of DNA methylation in sex hormones receptors in skeletal muscle, which could play into this hypothesis [18]. Altogether, this highlights the role of sex hormones in modulating miRNA expression in a sex-specific manner and potential role in programming sex-divergent targeting patterns.

Based on the sex-biased miRNAs, putative analysis revealed an enrichment of 7 TFs at baseline and 3 TFs in response to exercise including TGFβ. TGFβ and its downstream mediator SMAD3 regulate miRNA biogenesis [55, 56] and have an important role in skeletal muscle development [57, 58]. TGFβ putatively targeted 9 sex-biased miRNAs including mir-29a, which is inhibited by TGFβ and associated with altered substrate oxidation in skeletal muscle [59]. In the current study, miR-29a-3p expression was upregulated in response to the acute bout of exercise in males but not females. This could suggest that TGFβ and SMAD3 were inhibited with aerobic exercise in males but not females. There is limited literature that has conducted sex-stratified analysis in acute aerobic exercise studies although an acute bout of resistance exercise elicits a sex-specific response [60, 61]. Whether the aerobic exercise elicits a similar sex-specific TGFβ response has not been investigated. Altogether, our findings suggest that TFs may have sex-divergent targeting patterns of microRNAs in skeletal muscle in line with previous findings [16].

The miR-30 family is made of 5 miRNAs including miR-30a-5p, miR-30b-5p, miR-30c-5p and miR-30d-5p/-3p. The miR-30 family plays a regulatory role in the differentiation of human skeletal muscle cells by targeting genes associated with cell cycle, proliferation, and differentiation processes [42, 43]. In the current study, all 5 family members displayed differences in expression between sexes and a sex-specific response to exercise (supplementary fig 2). Specifically, miR-30a-5p and miR-30d-3p/-5p were upregulated after exercise in females but remained unchanged in males, while miR-30c-5p and miR-30b-5p were upregulated after exercise in males and remained unchanged in females. We therefore sought to investigate the effects of the miR-30 family on the myocyte transcriptome and to explore putative pathways in a sex-specific manner.

To investigate the extent of the maintenance of the sex-phenotype in primary cell lines, we analysed the baseline differences in the transcriptome between male and female donor muscle cells. A total of 27 genes were differentially expressed between the male and female donor cell lines. These included up-regulation of many Y-encoded (male-specific) genes that are considered to drive the sex differences at molecular level and included *Ddx3y* (DEAD box helicase 3), *Uty* (ubiquitously transcribed tetratricopeptide repeat gene on Y chromosome) and *KDM5D* (Lysine-specific demethylase 5D) amongst others (supplementary file 2) [62, 63]. Interestingly, the X-encoded paralogs (*Ddx3x, Utx, KDM5C*) were not different between the sexes, which may be due to X-chromosome inactivation ensuring dosage compensation across the sexes [64]. This finding adds to the existing evidence [65, 66] that sex differences originating from the sex chromosome complement seem to be conserved in primary cell lines [66]. Altogether, these data provide novel evidence that some sex-differences may be recapitulated in primary cell lines. However, the extent of the maintenance of sex differences mediated by gonadal sex hormones is still to be elucidated [62]

Before adjusting for the sex of the donor, initial analysis found differences in the transcriptome profile in response to overexpression of both miR-30c and miR-30a. In response to the overexpression of miR-30c, 108 genes were differentially altered, of which 55 of these genes have been experimentally validated to interact with miR-30c. MiR-30a overexpression induced changes in 247 genes, of which 123 of these genes have been experimentally validated to interact with miR-30a [34]. These genes broadly related to the biological pathways “protein glycosylation”, and “Golgi vesicle transport”. Protein glycosylation is an important post-translational modification and refers to the enzymatic process by which a carbohydrate binds, via covalent bonds to hydroxyl group of a protein[67]. Glycosylation is involved in many biological processes such as cell adhesion, receptor activation and signal transduction [67].

Glycosylated proteins play an important role in myogenesis and muscle development, with abnormal glycosylation associated with muscle disease and atrophy[68]. These results are consistent with models in other muscle tissues (cardiac and smooth), where the down-regulation of the miR-30 pathway results in muscle atrophy and its up-regulation promotes myogenesis and protein synthesis [43]. These data in a human primary cell-line provides preliminary evidence that the miRNA-30 family may play a role in myogenesis by targeting genes related to protein glycosylation.

Finally, we found that miR-30a/c target genes in sex-specific manner. This aligns with previous findings in cancer cell lines [69] and platelets[70], where other sex biased miRNAs have been found to target genes in a differential manner. For example, in response to overexpression of miRNA-30c *CASTOR2* (Cytosolic arginine sensor for mTORC1 subunit 2), a skeletal muscle enriched gene, was up-regulated in male donor cell lines only. CASTOR2 is a negative regulator of arginine signalling to the mTORC1 pathway and thus allows for increased cell proliferation [71]. In animal models, sex differences in the mTOR pathway signalling have previously been found [72, 73]. While it could not be determined whether miRNA-30c is directly targeting *CASTOR2* and therefore modulating cell proliferation via the mTORC1 pathway, putative evidence indicates with high confidence that miRNA-30c targets *CASTOR2* [74]. Further research is required to validate the role of miR-30a/c as upstream regulators of sex differences in skeletal muscle.

In conclusion, sex-differences in the expression profile of miRNAs were found in skeletal muscle at both baseline and in response to exercise. These miRNAs had regulatory roles in skeletal muscle development and our findings suggest miRNA play a profound but highly complex role in programming sex-differences in the skeletal muscle phenotype.

